# Liming pasture soils in the Amazon region promotes low-affinity methane oxidation by type I and II methanotrophs

**DOI:** 10.1101/2024.01.16.575838

**Authors:** Leandro Fonseca de Souza, Fernanda Mancini Nakamura, Marie Kroeger, Dasiel Obregon Alvarez, Moacir Tuzzin de Moraes, Mariana Gomes Vicente, Marcelo Zacharias Moreira, Vivian Helena Pellizari, Siu Mui Tsai, Klaus Nüsslein

## Abstract

In the Amazon Forest region, cattle pastures are the main land use subsequent to deforestation. This land-use change affects the soil microbial community and methane fluxes, shifting the soil from a methane sink to a source. Soil physical and chemical attributes are changed due to slash-and-burn processes, including an increased soil pH after forest-to-pasture conversion. Without amendments, the pasture soils can become acidic again resulting in many cases in soil degradation. Liming is a standard management practice to increase soil pH while decreasing Al^3+^ availability. Liming is important to recover these degraded lands and increase soil fertility, but its impact on soil methane cycling in tropical soils is unknown. Here we investigated the role of soil pH on methane uptake under high concentrations of the gas. The top layer of forest (pH 4.1) and adjacent pasture soils (pH 4.8) from the Eastern Amazon were subjected to liming treatment (final pH 5.8) and incubated with ∼10,000 ppm of ^13^CH_4_ for 24 days to label DNA with ^13^C. Soil DNA was evaluated with Stable Isotopic Probing (SIP-DNA), methanotrophic abundance was quantified (*pmoA* gene), and high throughput sequencing of 16S *rRNA* was performed. Liming increased the methane uptake in both forest (∼10%) and pasture (∼25%) soils. Methanotrophs *Methylocaldum sp*. (type I) and Beijerinckaceae (type II) were identified to actively incorporate carbon from methane in limed pasture soils. In limed forest soils, *Nitrososphaeraceae*, *Lysobacter* sp., and *Acidothermus sp*. were identified as ^13^C-enriched taxa. The enrichment of the archaeal family *Nitrososphaeraceae*, known as ammonia oxidizers, is correlated with an increase of ammonia monooxygenase genes, which code for an enzyme complex with wide substrate specificity that can also perform methane oxidation. In conclusion, liming Amazonian pasture soils not only contributes to the fertility and recovery of degraded areas but also has the potential to improve the oxidation of methane at high concentrations of this gas.

## Introduction

Methane gas (CH_4_) is 86 times more potent than carbon dioxide in retaining heat in the atmosphere over a 20-year horizon (IPCC, 2013), and its emissions to the atmosphere are mainly linked to human activities such as irrigated agriculture, decomposition in landfills, and livestock (IPCC, 2013). Tropical forest soils act as methane drains in the global methane (CH_4_) cycling process (Keller et al., 1986). However, deforestation followed by the establishment of pastures, converts these soils to sources of atmospheric methane (Fernandes et al., 2002; Fonseca de Souza et al., 2022; Goreau & de Mello, 1988; Meyer et al., 2020; Steudler et al., 1996; Venturini et al., 2022). The process of establishing pastures in Amazon Forest areas is commonly preceded by selective logging and vegetation burning, which leads to the incorporation of alkaline ashes into the soil promoting an increase in pH (Fearnside & Barbosa, 1998; Fritze et al., 1994). In the years following pasture establishment, there is a tendency for the soil to acidify again, reaching pH values close to those observed in forest areas (de Moraes et al., 1996). The liming practice allows for acidity correction and calcium and magnesium supplementation of the soil; however, it can also impact methane cycling (Kunhikrishnan et al., 2016; Zhang et al., 2022).

Atmospheric methane is produced mainly from biotic processes (70-80%). Methanogenic archaea are the major known source, but 5% of it can be biologically removed from the atmosphere by methanotrophic microorganisms, which also prevents methane from being emitted from soil (Canadell et al. 2021). Methanotrophic bacteria are ubiquitous and found in all types of soils, oceans, and extreme environments (Conrad, 2009; Kussmaul et al., 1998). They are Gram-negative and classified in the phylum Proteobacteria (syn. Pseudomonadota), in the Gammaproteobacteria (type I) and Alphaproteobacteria (type II) classes, which harbor the most abundant and well-known methanotrophs (Knief, 2015; Park & Lee, 2013; Semrau et al., 2010), as well as in the Verrucomicrobia (syn. Verrucomicrobiota) (type III) (Op den Camp et al., 2009). Archaea can also oxidize methane by anaerobic pathways, but this is more common in aquatic environments (Segarra et al. 2015). In bacteria, the initial stage of incorporation of methane is through its conversion to methanol. This step is mediated by the enzyme methane mono-oxygenase, which is encoded by conserved genes that have been used as a marker for studies of methane oxidation in environmental microbiology: *pmoA* (constituent of the beta subunit of pMMO) and *mmoX* (constituent of the alpha subunit of sMMO) (Hanson & Hanson, 1996; Knief, 2015). The separation into major types (I and II) is based on the biochemical pathway used to assimilate the carbon from formaldehyde, formed after methane conversion to methanol. Type I methanotrophs are Gammaproteobacteria that assimilate formaldehyde via the ribulose mono-phosphate pathway, while type II are Alphaproteobacteria that use the serine pathway. Typically, type I and type II methanotrophs can incorporate methane under high concentrations of methane gas (low affinity) while type II are also able to act under low concentrations (high affinity) (Cai et al., 2016; Dunfield et al., 1999).

The conversion of forest to pasture in the Amazon region is a main driver of changes to soil bacterial community structure and diversity (de Carvalho et al., 2016; Jesus et al., 2009; Rodrigues et al., 2013). Previous studies investigating the responses of methanotrophs and methanogens to forest-to-pasture conversion observed a reduction in the relative abundance of methanotrophs (16S *rRNA*), as well as in the methanotroph marker genes (*pmoA*), and an increase in the main molecular marker for methanogenic archaea (*mcrA*) (Meyer et al., 2017; Paula et al., 2014). More recently, Kroeger et al. (2021) demonstrated that forest-to-pasture conversion not only increased the abundance of methanogens in pastures, but also stimulated their activity, and the higher soil pH in pastures was suggested as one of the drivers of this change. Although, the optimum pH of most cultivable methanotrophs is neutral (between 6.8-7.0) (Whittenbury et al., 1970), the wide pH range where methane oxidation occurs in forests suggests the presence of a diverse community acting in the process (Saari et al., 2004). In temperate forest soils, the effect of pH on methanotrophic distribution is decisive, where Gammaproteobacteria are more frequent in soils with pH higher than 6,0, and Alphaproteobacteria are likely to be more frequent in acidic soils (Knief et al., 2003).

Microbial activity is an important property for understanding biogeochemical processes, but it is not easy to access given the complexity of interactions between microorganisms in the environment. Using Stable Isotope Probing (SIP), it is possible to evaluate the active fraction of microorganisms by enriching their nucleic acids after growing with substrates enriched with heavy isotopes. A subsequent density fractionation allows the identification of those microorganisms within the community that actively incorporated a certain substrate to make it part of their biomass (Radajewski et al., 2000). Since DNA-SIP relies on cellular replication to obtain the adequate signal of the labeled substrate in the active microorganisms, it is often necessary to subject the samples to high concentrations of the substrate (Neufeld et al., 2007). These concentrations should ideally be close to concentrations to which the microorganisms are exposed in their natural environment. Analyzing the metabolism of methane in soil brings added complexity since the soil environment is heterogeneous and may be under concentrations ranging from 1.8 ppm (atmospheric) to 14,000 ppm (landfills) (Gebert & Perner, 2015). Pasture soils in the Eastern Amazon can emit up to 6 µg of C-CH_4_ per gram of soil if incubated at saturation (100% water holding capacity) for 30 days, equivalent to 46,667 ppm of CH_4_ (Venturini et al., 2022). These results indicate a high potential for methane production by tropical soils under saturated conditions. Knowing that methane oxidation in nature is used by groups of microorganisms that have rather different affinities for methane, methane concentration is an important variable to be considered, since incubation itself can favor certain groups. In this study, we hypothesized that soil pH correction to a pH value preferred by known methanotrophs would positively stimulate the activity of methanotrophic microorganisms. We tested acidic forest and pasture soils in the eastern Amazon rainforest to evaluate the impact of soil liming on methane uptake by low-affinity methanotrophs. The composition of soil microbial communities was determined by high throughput sequencing of 16S *rRNA* genes and combined with activity measurements of soil gas fluxes and SIP. This combination of analyses enabled us to identify which soil microorganisms more actively consumed methane following soil pH correction by liming.

## Materials and Methods

### Sampling sites

Studies were performed with soils from Eastern Amazonia, from the National Forest of Tapajós, and nearby farms of Belterra, state of Para, Brazil (52° 25′48″ S 54° 43′12″ W). Soils were sampled in October 2017 (Supplementary Table S1) and April 2018, in both dry and rainy seasons, respectively. The Tapajós National Forest is a conservation area. The adjacent pasture was established between 1989-1994, and is characterized by sparse signs of degradation, no history of liming, and burned irregularly to control invasive plants. Cattle were present at sampling time. In each site, a square plot of 100×100m (1 ha) was selected and five sampling points were chosen located in the corner and center of this square, and 20–30 kg of soil from the upper 0–10 cm layer were sampled from each point. The soil samples were conserved in plastic bags that had a small opening protected by filter paper to ensure air exchange while minimizing soil water loss. The soil samples were transported by car to the Cell and Molecular Biology Laboratory at the University of Sao Paulo, CENA-USP, where sub-samples were used for chemical analysis, and the remaining soil was conserved at ambient temperature in the shade until the experiment started.

### Characterization of the chemical properties of soil samples

Chemical analyses of soil were performed at the Chemical Analysis Laboratory of Luiz de Queiroz College of Agriculture (ESALQ/ USP), using protocols by van Raij et al. (2001). The measured soil attributes were: pH in CaCl_2_; available phosphorus (P), potassium (K), calcium (Ca), and magnesium (Mg), by extraction in ion exchange resin; aluminum (Al) by extraction of potassium chloride 1 mol.L^-1^; potential acidity (H+Al), by SMP methodology (Shoemaker et al. 1961); organic matter by the dichromate titrometric method; boron (B) by extraction with hot water; copper (Cu), iron (Fe), manganese (Mn) and zinc (Zn) extracted by the DTPA-TEA extractor (pH 7.3); additional results were the calculations of total bases (BS), the potential Cation Exchange Capacity (CEC_pH 7_), base saturation (V%), and aluminum saturation (m%).

### Soil liming

The liming experiment was established with soils collected during the 2018 sampling event in a research greenhouse at the Center for Nuclear Energy in Agriculture - University of São Paulo (22°42’27.7"S, 47°38’41.0"W). The experimental setup evaluated the factors of land use (forest vs. pasture) and liming (with or without) of two soils as follows: the five sampling points from each 100×100 m area were combined, homogenized, sieved (5 mm), and placed in 10 L pots with approximately 5 kg of soil each forming a 10 cm layer. Each treatment had four replicates. The treatments were: – Forest soil at natural pH, Forest limed soil, Pasture soil at natural pH, and Pasture limed soil. The liming was performed by the addition of CaCO_3_ powder (98% purity, Synth laboratories, Diadema, SP, Brazil), with a Total Relative Neutralizing Power (TRNP) equivalent of ∼100% so that the samples reached a final pH of ∼6.0 (water), with a calculated Base Saturation of 65-75%. The moisture of all soils was standardized at the beginning of the experiment to ∼70% of the water holding capacity and readjusted every two to four days taking as reference the weight variation after drying 5 g of soil for 48 h at 105°C. Forty days after liming (reaction time to liming stabilization), the soil was sampled to prepare the incubation with ^13^CH_4_.

### ^13^CH4 incubation and DNA fractionation

Based on the results of preliminary soil incubations with ^12^CH_4_ at a range of concentrations, we chose a concentration of 10,000 ppm for the incubation with ^13^CH_4_ gas (Supplementary Material). The microcosms were prepared in airtight 120 mL glass vials with 10 g of soil each, taken directly from the pots in the greenhouse. The atmospheres inside the microcosm vials were adjusted by the addition of 1.4 mL of ^13^CH_4_ (99% purity - Cambridge Isotope Laboratories, Tewksbury, MA, USA) or ^12^CH_4_ (99.5% purity - White Martins, Piracicaba, SP, Brazil) after the vials had been sealed with butyl rubber stoppers. To each vial, 20 mL of synthetic air (O_2_ and N_2_ in atmospheric concentrations - White Martins, Piracicaba, SP, Brazil) were added to generate an excess of 20% of internal pressure to allow detection of possible gas leakage. These incubations were carried out with five technical replicates for two soils and two pH values: Forest-natural pH, Forest-liming, Pasture-natural pH, and Pasture-liming, with either ^13^CH_4_ or ^12^CH_4_ (5 replicates x 4 treatments x 2 methane isotopes). Over 24 days, methane concentrations were measured every 3-4 days, followed by venting and replacement of the methane-enriched atmosphere. All replicates were evaluated with gas measurements, but soils from three vials of each treatment were randomly selected for molecular analysis.

The application of 1.4 mL of ^13^CH_4_ every 3-4 days would allow the approximate incorporation of 50 μmol of ^13^C into the biomass in 10 grams of soil, a minimum recommendation by Neufeld et al. (2007). As the efficiency of incorporation likely varies among treatments, 6 cycles of renewal of the atmosphere were performed over the 24 days of the incubation experiment.

The incorporation of ^13^C isotopes from ^13^CH_4_ into the microbial biomass was determined at two-time intervals of the experiment, after 14 and 24 days. The isotopic ratio of carbon (δ^13^C) in the respired carbon dioxide (CO_2_) was quantified using a pre-concentrator (PreCon, Thermo Fisher Scientific, Waltham, MA, USA) coupled to an isotopic ratio mass spectrometer (Delta Plus Model, Thermo Fisher Scientific, Waltham, MA, USA) with a continuous flow. A sub-sample of the gas (50-300 μL, depending on the concentration of CO_2_), was injected into the PreCon where the carrier gas (Ultra Pure Helium) led the sample to capillaries immersed in liquid nitrogen to be cryogenically concentrated. The sample was then pushed into a 25 m chromatographic column for the separation of CO_2_ under 20 psi of He (Ehleringer & Cook, 1998). The isotopic composition of carbon dioxide (δ^13^C-CO_2_) was expressed in deviation per thousand (‰) in relation to international standards V-PDB (Vienna PeeDee Belemnite) and the enriched concentration calculated from the total CO_2_ concentration, previously measured in gas chromatography.

### CH_4_ concentration measurements

To evaluate the concentrations of CH_4_ in the microcosm atmosphere, samples of 5 mL of gas were extracted from the headspace and injected in a gas chromatograph GC SRI 8610C (SRI Instruments, Torrance, CA) equipped with a Flame Ionization Detector. The detector was previously calibrated with reference gases (White Martins, Piracicaba, SP, Brazil) depending on the sample concentrations. The gas concentration was determined by heating at 250°C at a pressure of 143 kPa, with N_2_ as a carrier gas, an H_2_ pressure of 75 kPa, an air pressure of 50 kPa, and a retention time of 2.5 minutes.

### Extraction of DNA

The total DNA from the soil was extracted with the DNeasy PowerMax Soil Kit (Qiagen, Hilden, Germany) from 10 g of soil, according to the protocol provided by the manufacturer. The quantity and quality of the DNA samples were analyzed using a Qubit 4.0 fluorometer (Thermo Fisher Scientific, Waltham, MA, USA) and a Nanodrop 2000c spectrophotometer (Thermo Fisher Scientific, Waltham, MA, USA). The total DNA extracted from the samples was stored at -20 °C.

### Fractionation of DNA by ultracentrifugation

Total DNA was exposed to ultracentrifugation in a density gradient of cesium chloride (1.725 g/ mL), and subsequently fractionated with the method adapted from Neufeld et al. (2007). The ultracentrifugation vials were quick-seal polypropylene 2.0 mL 11×32 mm tubes from Beckmann Coulter (Part. 344625, Brea, CA, USA). The tubes were loaded with 1 μg of DNA in 20 μL of water (50 ng/μL), filled with CsCl solution, sealed as recommended by the manufacturer, and then centrifugated at 64,000 rpm and 20°C for 40 h in a Beckman Optima TLX ultracentrifuge with a TLA-120.2 rotor (Brea, CA, USA). The fractionation of the density gradient was performed with the aid of a peristaltic pump, pressing a syringe with a solution of resazurine (0.1%) connected by a hose with a needle at the terminal to the top of the tubes, generating 20 fractions of 100 μL from a hole on the bottom of the tube by an output of 3.3 μL/s. The density gradient formation was evaluated using a refractometer (Reichert AR200; Depew, NY, USA). The DNA was precipitated and purified by washing with 2 volumes of polyethylene glycol (6000) and 1 volume of ethanol (70%), following the protocol of Neufeld et al. (2007), except for the use of 1 μL of linear acrylamide (5 mg/mL) instead of glycogen in the DNA precipitation stage. Total DNA concentrations per fraction were measured using a Qubit 4.0 fluorometer (Thermo Fisher Scientific, Waltham, MA, USA).

### Quantification of the methane consumption marker gene (qPCR)

Quantitative PCR (qPCR) technique was used to quantify the number of copies of the *pmo*A gene from total soil DNA samples, using primers A189F (5’-GGNGACTGGGACTTCTGG-3’), as in Holmes et al. (1995) and MB661R (5΄-CCGGMGCAACGTCYTTACC-3΄), in Costello & Lidstrom (1999). The reaction conditions were 95°C for 10 min followed by 45 cycles of: 95°C for 30 s, 58°C for 30 s, and 72°C for 45 s, followed by a plate read, incubation at 68°C for 5 min, melting curve from 65°C to 95°C with a read every 1°C, and temperature holds of 1 s. A standard curve was built in five steps from 10^2^ to 10^5^ copies of a target gene previously obtained by PCR from a pure culture of *Methylosinus sporium* (DSMZ 17706). The qPCR was performed in triplicate for each soil DNA sample in a StepOne Plus (Thermo Fisher Scientific, Waltham, MA, USA), with a final volume of 10 μL, containing 5 μL of SYBR Green ROX qPCR (Thermo Fisher Scientific, MA, USA), 1 μL of each primer (5 pmols), 1 μL of DNA (adjusted previously to 10 ng/μL), 0.8 μL of bovine albumin (20 mg/mL) (Sigma-Aldrich, San Luis, MO, USA), and 1.2 μL of ultrapure water (Milli-Q).

Bias in the analysis of results between plates and runs was minimized by quantifying genes using the LinRegPCR software (Ramakers et al., 2003), in which the raw amplification data of each sample was used to calculate individual reaction efficiencies, and thresholds were established for each group of technical replicates. The data generated in arbitrary fluorescence units were converted to the number of copies of the genes using linear interpolation between the known quantities in the standard curve (5 points) and the observed fluorescence measurements, using the curves of each plate as a reference for the respective samples.

### Sequencing of 16S *rRNA* gene fragments

To access the composition of the microbial community, we used high throughput sequencing technology in a MiSeq Illumina platform (Illumina, San Diego, CA, USA). We used the MiSeq V3 600c paired-end sequencing kit with the primers 515F (5’-GTGYCAGCMGCCGCGGTAA-3’) (Parada et al., 2016) and 806R (5’-GGACTACNVGGGTWTCTAAT-3’) (Apprill et al., 2015) for the V4 region of the 16S *rRNA* gene (Caporaso et al., 2011) at the Functional Genomics Center of the Luiz de Queiroz College of Agriculture (USP). This kit was chosen for the number of reads generated, the size of the quality paired-end reads, and considering the diverse environment that soil represents.

The gene library was prepared as follows. PCR reactions were performed with 2.0 μL of DNA (10 ng/μL), 0.5 μL each of Forward and Reverse Primers (10 μM), 12.5 μL of 2x PCR Bio Ultra Mix (PCRBIO, Wayne, PA, EUA), and 9.5 μL of PCR grade water in a total volume of 25 μL. The reaction conditions were: 94°C for 3 min followed by 30 cycles of 94°C for 30 s, 50°C for 30 s, 72°C for 30 s, and a final extension at 72°C for 10 min. After the PCR, DNA purification was performed using AMPure XP beads (Beckman Coulter, Brea, CA, USA) and agarose gel verification. Nextera XT adaptors were used (Illumina, San Diego, CA, USA), followed by another purification with AMPure XP beads and verification in agarose gel. The amplicon pools for each primer were made from equal volumes using quantification by qPCR with the KAPA Illumina quantification kit (Roche, Basel, Switzerland).

Demultiplexing of the data was performed using the Illumina tool bcl2fastq version 2.20 and data computational processing was performed using the pipeline QIIME2 (vs. 2019.10) (Bolyen et al., 2019). The primers were trimmed from reads with cutadapt plugin (Martin, 2011), and data quality control was achieved using the DADA2 tool (Callahan et al., 2017), without clustering into OTUs (Operational Taxonomic Units). Forward and reverse sequences were truncated for DADA2 analysis based on quality scores at positions 280 and 250, respectively. Taxonomic identification was assigned using the q2-feature-classifier tool (Bokulich et al., 2018) with the SILVA database (vs. 132) at a cutoff of 99% (Quast et al., 2013), trained for the primer set.

Sequencing and supporting data are available in Zenodo, on https://doi.org/10.5281/zenodo.8144870.

### Phylogenetic analyses

Phylogenetic trees were generated based on 16S *rRNA* gene fragments. The sequences were aligned and grouped into trees using CLC Genomics Workbench 20.0 software (QIAGEN, Aarhus, Denmark) with default parameters, and the maximum likelihood model (PHYML function) with Unweighted Pair Group Method with Arithmetic mean (UPGMA) assuming even replacement frequencies to the bases (Kimura, 1980). Results were tested by simulation with 1,000 bootstrap repeats. The reference sequences of the Beijerinckiaceae family were obtained from the curated database RDP version 11 (Cole et al., 2014), only sequences greater than 1,200 bp of good quality and of type strains, and truncated according to the primer set. The only sequence of the 16S *rRNA* gene available for a methanotrophic bacterium USCα was also used (Pratscher et al., 2018).

### Statistical analysis

For the analysis of significant differences within the qPCR and CH_4_ consumption data, we applied the Kruskal-Wallis test when the data were not normally distributed, or when data were normally distributed the analysis of variance (ANOVA) with post hoc Tukey HSD test (Honestly Significant Difference) using the agricultural package, version 1.2-8, in the software R Studio (R Core). For the data resulting from sequencing, in the QIIME2 2019.10 environment, the ANCOM tool was used, which compares the relative abundance of each ASV (Amplicon Sequence Variant) or rate between two groups per logarithmic reasons (Mandal et al., 2015). ANCOM generates volcano plots, which display the centralized logarithmic ratio on the X-axis of the graph, and on the Y-axis a dimensionless indicator W which represents the number of times the null hypothesis (the average abundance of an ASV in one group is equal to that of another group) was rejected for a given ASV.

### Differential abundance analysis between bacterial groups

Two conditional restrictions had to be established to identify microorganisms that have incorporated ^13^C from ^13^CH_4_ into their DNA while avoiding those with increased relative abundance in the high-density DNA region for a reason other than ^13^C incorporation (Supplemental Figure S4). The conditions were I) the ASV sequence should be enriched in the high-density region of the ^13^CH_4_ incubation in relation to the high-density region of the ^12^CH_4_ incubation. This condition allows the detection of microorganisms that incorporated ^13^C into their DNA despite its buoyant density related to their DNA GC content. And, II) the ASV should not be enriched within the high-density region of the ^12^CH_4_ incubation, compared to the lighter-density region. Otherwise, the enrichment of this ASV could be due to other factors such as GC content in the sequence and not incorporation of ^13^C. Only those ASVs that fit both conditions were recognized as representative of active microorganisms incorporating ^13^C from ^13^CH_4_.

The condition of enrichment in the high-density fraction in relation to the low-density fraction in the ^13^CH_4_ incubation was applied as a complementary criterion. This decision was based on the fact that it is not feasible, through general DNA analysis, to separate ASVs from living, dormant, or dead cells. The incubation lasted 24 days, and it would be possible to have more DNA from dormant/dead methanotroph cells in the light region than in the heavier fractions, enriched in ^13^C, which could generate a false negative result of non-incorporation activity of ^13^C.

We evaluated the quality of the gradient formation by accounting for the GC content in genomes from microorganisms enriched in the high and low-density fractions. ASVs significantly enriched in the light and heavy fractions were grouped into their categories. Then the genomes completely sequenced, with phylogenetic affiliation as close as possible to each ASV, were consulted in the genome database of the NCBI (National Center for Biotechnology Information). The GC content information of these reference genomes was used as a representative for the identified ASVs to determine the differences between the low-vs. high-density sets by ANOVA (Tukey HSD post hoc).

## Results

Soil preparation started 40 days before incubation experiments when all soil samples were potted and stored in a greenhouse. Half of the soil was amended with CaCO_3_ to control for natural acidity. Liming showed satisfactory results, with a reduction in potential acidity, in total available aluminum, and an increase in calcium and soil pH (Supplemental Table S1).

The appropriate concentration of CH_4_ added in the incubation of our SIP study was evaluated in a preliminary experiment with different concentrations of ^12^CH_4_ in pasture and forest soils (Supplemental Figure S1). The results indicated a concentration of 10,000 ppm of ^13^CH_4_ for the SIP incubation, which was high enough to observe the metabolic incorporation of ^13^C into DNA after 24 days.

The soil demonstrated an increasing capacity for methane uptake over time in all treatments, however, the uptake was significantly higher in the limed soils (Figure 1). Pasture soils consumed methane at much higher rates, especially pH-adjusted pasture soils where added methane was almost completely consumed after the first nine days. Except for limed pasture soils, methane consumption decreased between days 9 and 13 for most soils.

**Figure 1.**
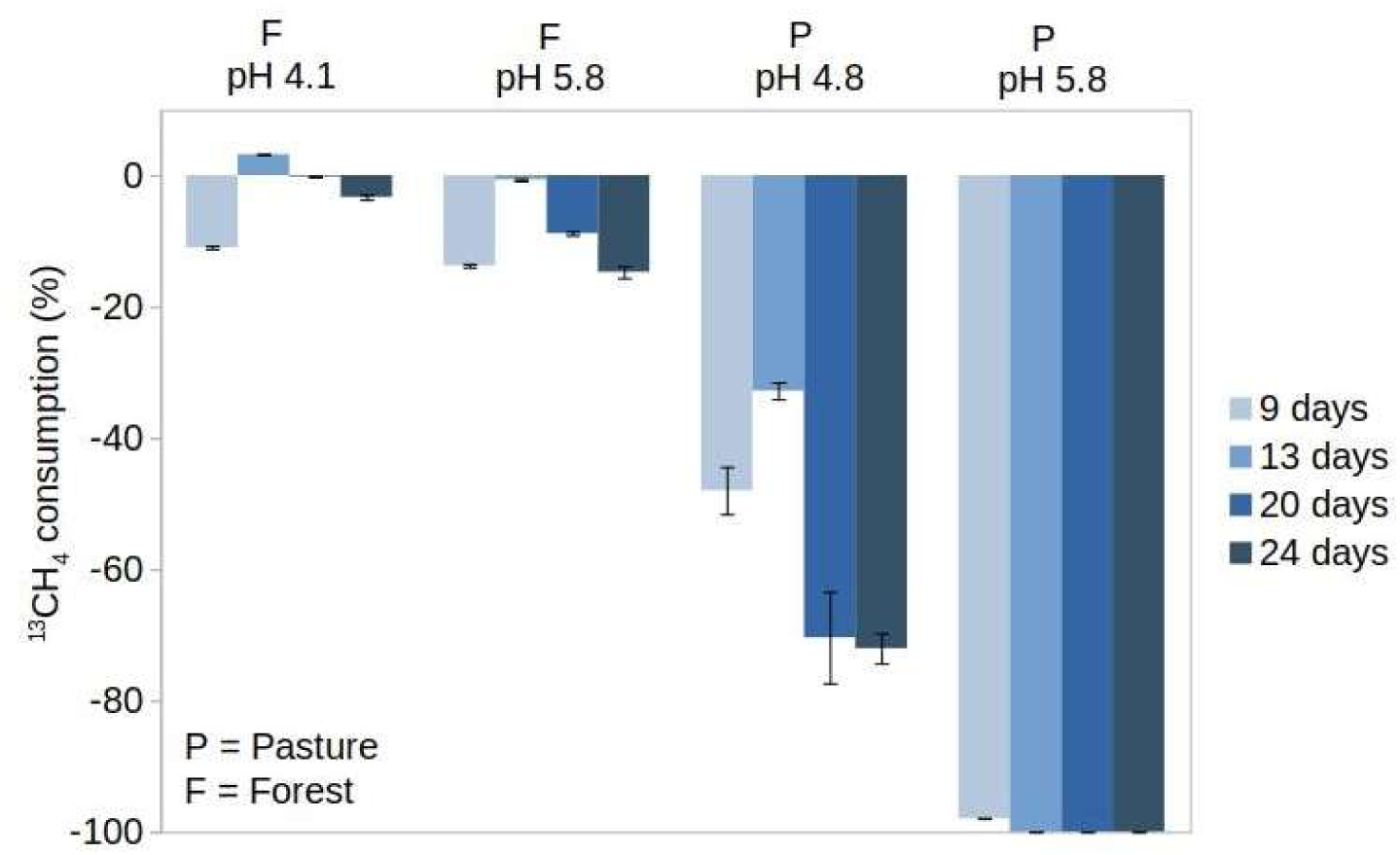
Percent of ^13^CH_4_ removal (initial ∼10,000 ppm) by forest and pasture soils with and without liming after 3-4 day long incubation cycles across 24 days of incubation. Data collection started at day 5, to allow microcosm to stabilize. No data is available for day 17. Bars indicate standard deviation.

To estimate how much of the ^13^C from ^13^CH_4_ was incorporated into the microbial biomass, the ratio of ^13^CO_2_/ ^12^CO_2_ in the microcosm headspace was evaluated and analyzed together with the ^13^CH_4_ concentration data (Tveit et al. 2019). There were no differences in the ^13^CO_2_ concentrations between treatments with and without liming for each soil (Figure 2), despite the differences in methane oxidation (Figure 1). This indicates possible differences in the methanotrophic metabolism following soil liming, with higher storage of the carbon from ^13^CH_4_ and cell growth of methanotrophs in limed soils, consuming more ^13^CH_4_ and releasing ^13^CO_2_. The greatest incorporation of ^13^C occurred in the pasture-limed soils, followed by the pasture soil with natural pH and limed forest soil. Less than 5% of ^13^C was incorporated into biomass in acidic forest soil, compared to more than 70% in limed pasture soil (Figure 2).

**Figure 2.**
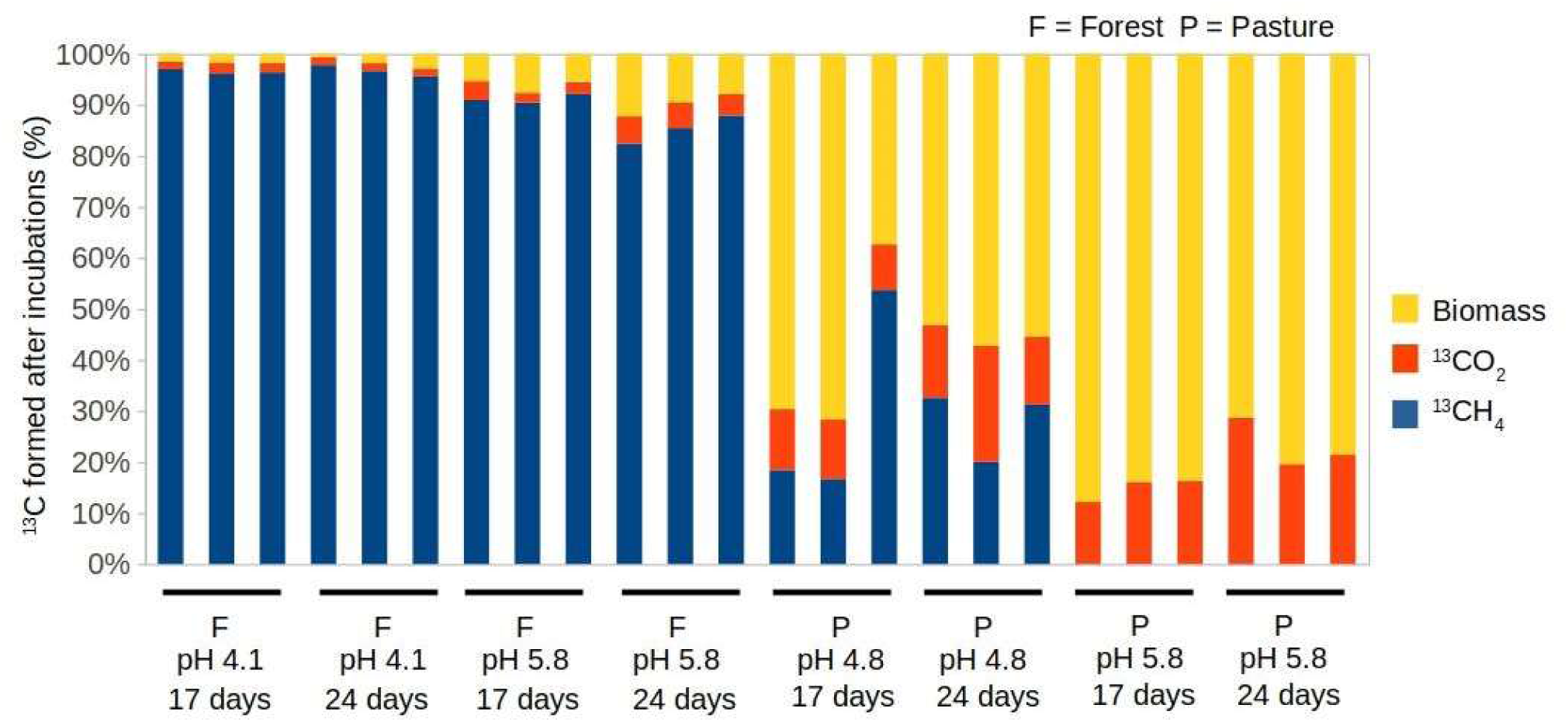
Relative incorporation of ^13^C into soil microbial biomass. Incorporation was estimated based on the initial concentration of ^13^CH_4_ (100%) in the microcosms from which the remaining concentrations of ^13^CH_4_ and ^13^CO_2_ after 4 days were subtracted. Data are shown for 17 and 24 days after the incubation started.

After estimating the amount of ^13^C incorporation into the microbial biomass (Figure 2), total soil DNA was extracted and analyzed by SIP. The fractionation of DNA resulted in three regions, named hereafter Very High (VH), High (H), and Low (L) density (Supplemental Figure S2). Fraction number 11 was defined as the divisor of the low-density fractions and, when enough DNA was present, a second division between high and very high-density was added in fraction 6. The VH fractions were obtained only for pasture soils (Supplemental Figure S2).

As an indicator of appropriate gradient formation, we estimated the genomic GC %mol content of the microorganisms significantly enriched in each group of fractions (L and H). Significant differences were observed, with genomes of high GC %mol content microorganisms enriched in higher density fractions by an average of 15% (Figure 3).

**Figure 3.**
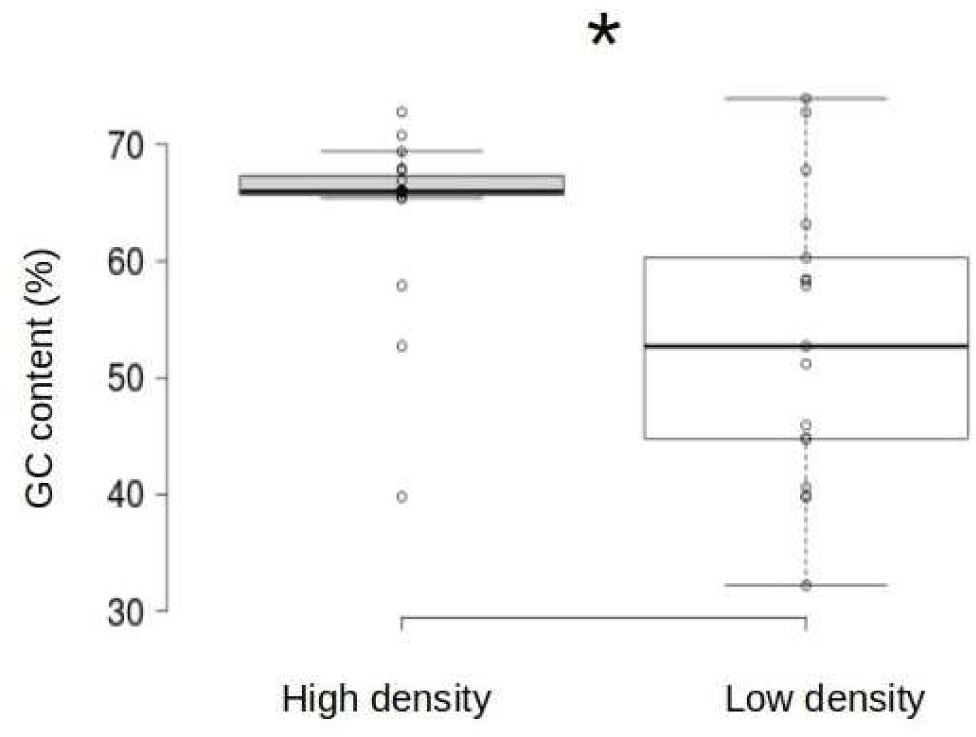
Differential abundance of representative genomes of the sequences (ASVs) in High-density and Low-density regions based on their GC %mol content using ANCOM. Data for all treatments are included (Forest and Pasture with and without liming). * Kruskal-Wallis (H=7.78; p = 0.005).

To estimate if liming stimulated the growth of methanotrophs, the abundance of methanotrophic bacteria was assessed by quantifying the number of copies of the *pmo*A gene in the H and L fractions of the ^12^CH_4_ and ^13^CH_4_ incubations. No significant enrichment was observed in forest soils with or without liming. In pasture soils, however, enrichment was seen for both treatments with higher *pmo*A gene abundance in the limed pasture soils (Figure 4).

**Figure 4.**
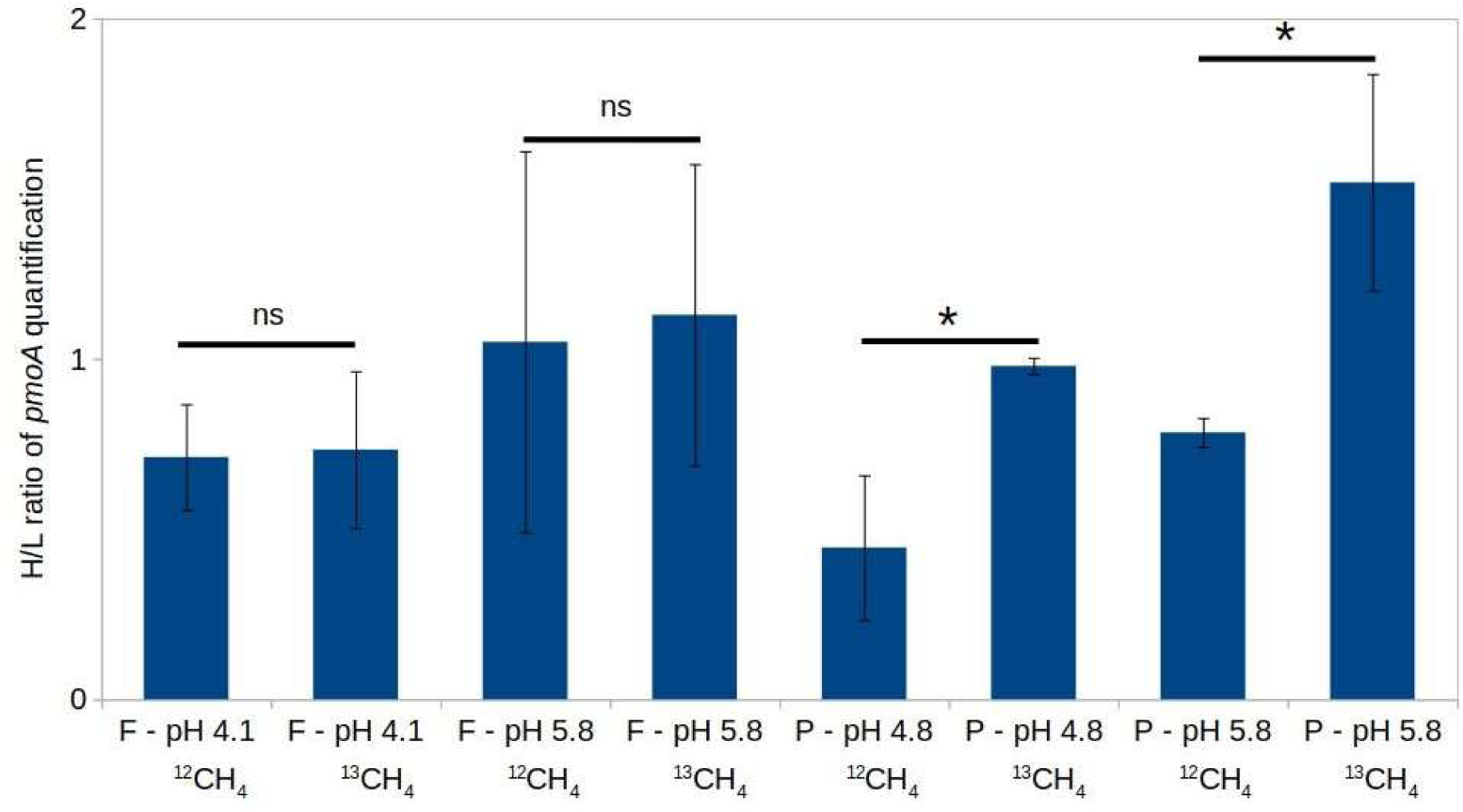
Ratio of the number of copies of the *pmo*A gene between the H (1-11) and L (12-20) fractions in the ^12^CH_4_ and ^13^CH_4_ incubations. *Indicates significant differences (Tukey HSD; p <0.05) and ns = not significant. F = Forest and P = Pasture. pH measurements in water.

After the enrichment of methanotrophic bacteria in the high-density fractions of ^13^CH_4_ incubations was confirmed, high-throughput sequencing of the 16S *rRNA* gene was carried out to identify these microorganisms. The results showed an adequate sequencing profile, with saturation of rarefaction curves below the minimum sequencing depth (Supplemental Table S2, Supplemental Figure S3). The analysis indicated active groups in the ^13^C incorporation from ^13^CH_4_, after comparing ASVs enriched in the high-density fractions of ^13^CH_4_ incubations (H13), with the dense region in the ^12^CH_4_ (H12) incubations. To ensure that these ASVs would not be enriched in the high-density fractions for reasons other than the incorporation of ^13^C, the taxonomic groups enriched in this analysis should not be enriched in the H12 region in relation to the L12 region (Supplemental Figure S4). No significant differences were observed (H13 x H12) in forest soils, or pasture soil without liming. However, enriched ASVs were detected in the limed pasture soil (H13 x H12), and were not enriched in the controls (L12 x H12). *Methylocaldum sp*. (type I methanotroph) was one of the microorganisms active in the incorporation of methane after liming pasture soils (Figures 6 and 7). Two other ASVs of microorganisms active in methane oxidation were identified as Beijerinckiaceae (type II methanotroph) (Figures 5 and 6).

**Figure 5.**
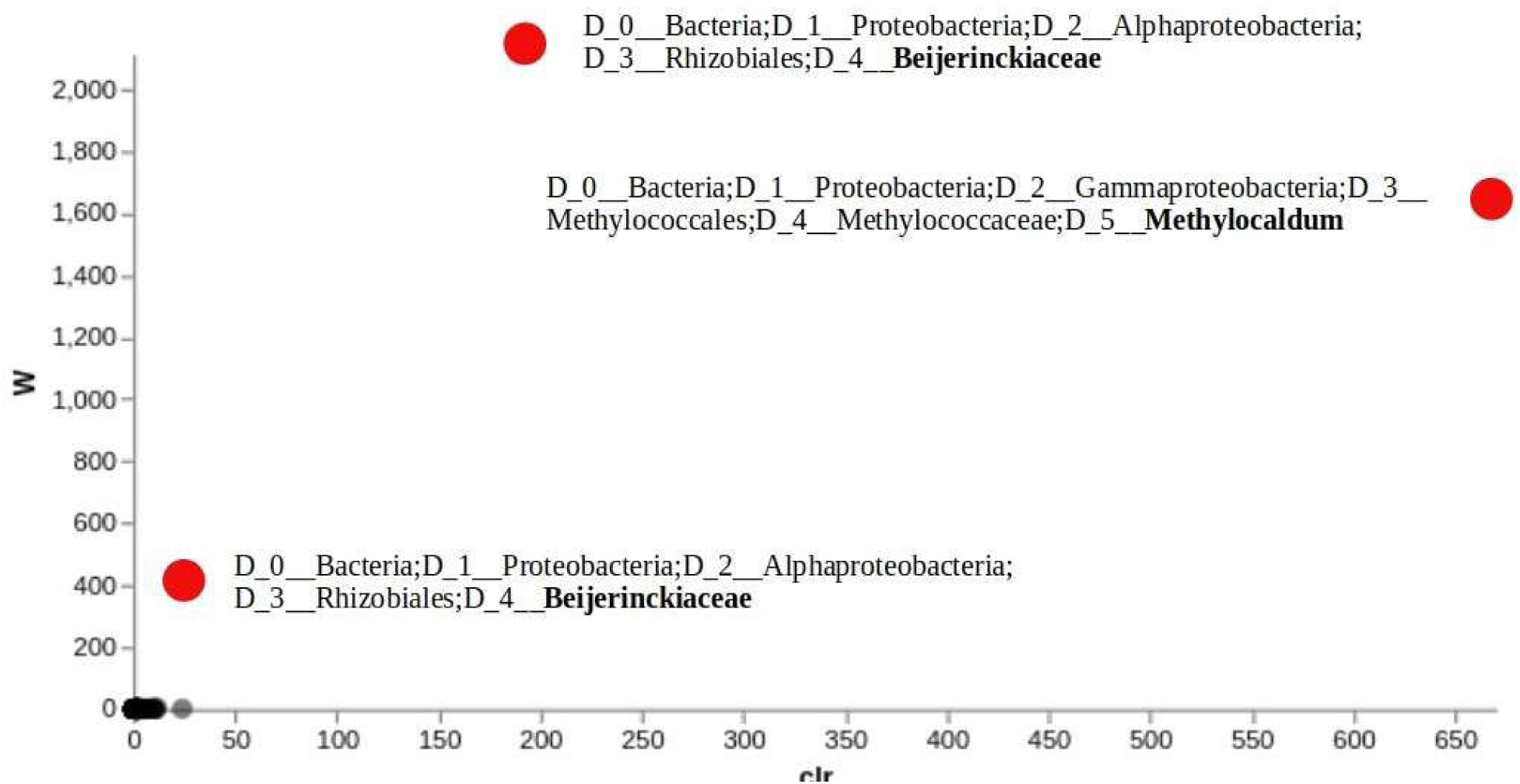
ASVs enriched in limed pasture soils (pH 5.8) in the Heavy (H) and Very Heavy (VH) fractions from SIP incubations with ^12^CH_4_ vs ^13^CH_4_. A calculation of the Centered Log Ratio (CLR) indicates how much a group is enriched in relation to the average relative abundance of a respective ASV, and W indicates the consistency of this observation about the other ASVs. The higher the values of W and CLR are, the greater and more consistent the difference in relative abundance. ASVs enriched in ^13^C are highlighted in red. All other ASVs are at the bottom of the graph, between CLR 0-50.

**Figure 6.**
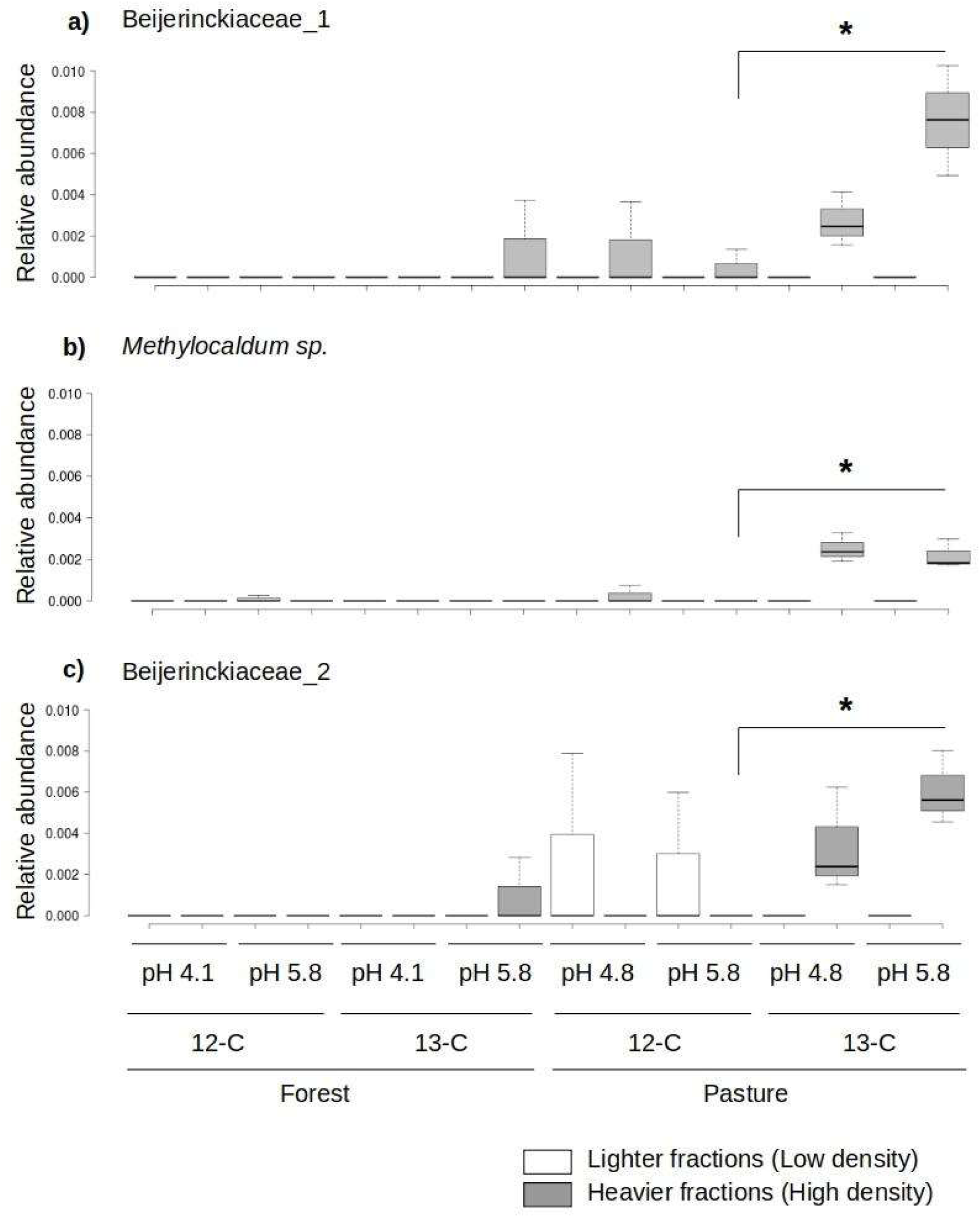
Relative abundance of ASVs from (a, c) Beijerinckiaceae *and (b) Methylocaldum* sp. between heavy fractions (H) from the ^12^CH_4_ and ^13^CH_4_ incubations. *ANCOM significant.

**Figure 7.**
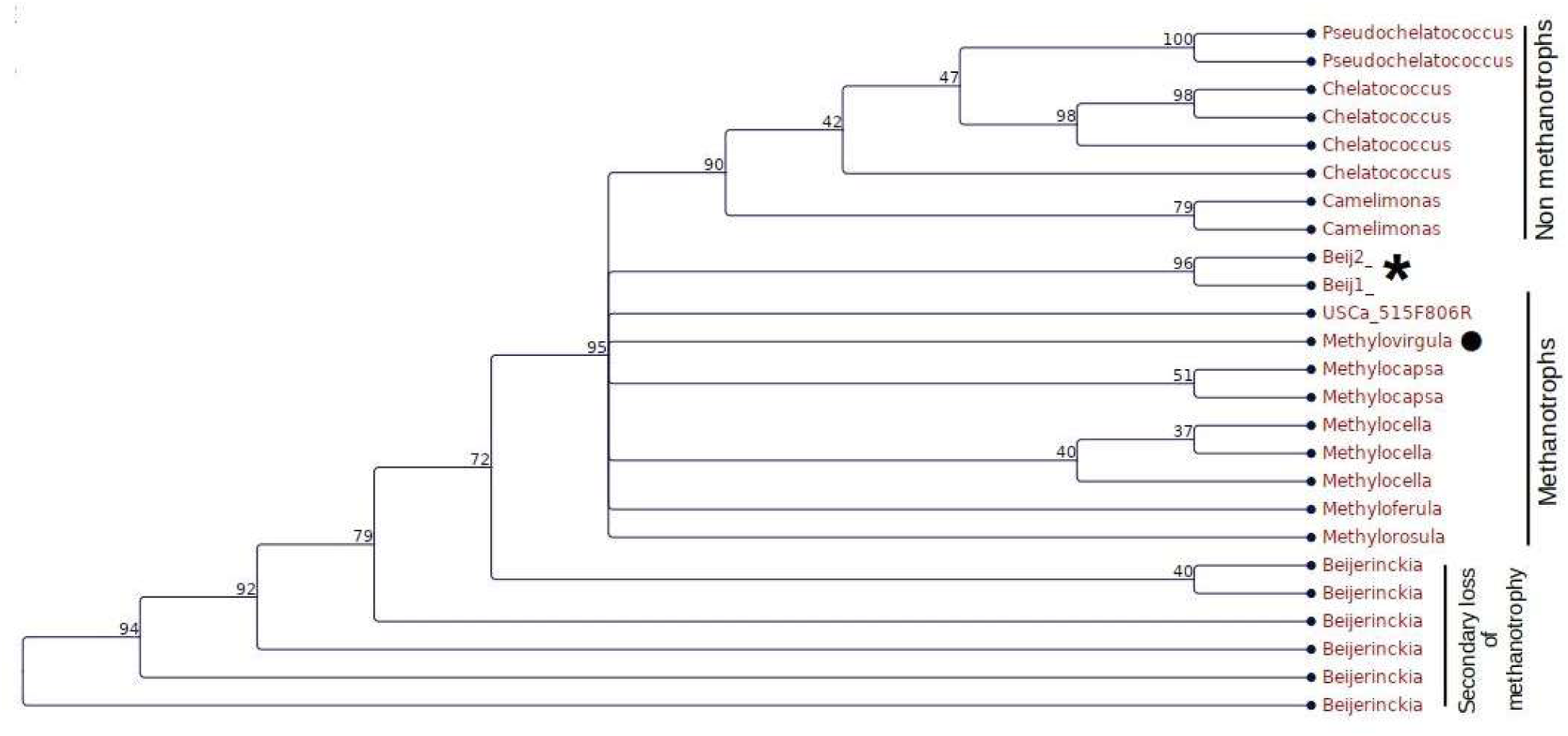
Cladogram built with sequence fragments of the 16S *rRNA* gene (V4 region) of ASVs identified in this study (* = Beij1 and Beij2) and reference sequences for the Beijerinckiaceae family obtained from the database RDP version 11. Numbers at nodes indicate bootstrap values after 1000 simulations. ● = not a methanotroph, but a facultative methylotroph.

The phylogeny of ASVs identified as Beijerinckiaceae was analyzed alongside with high-quality sequences available in a cured database for this bacterial family. The two ASVs are closely related, possibly the same genus, and cluster together with other methanotrophic genera within the family (Figure 7).

Finally, the H13 x L13 regions were compared for limed forest soils to provide additional information about the active microbes. Active microorganisms will not be detected when the abundance of their DNA falls below the abundance of inert or dead cell DNA already existing in the soil. The results show three taxa with direct or indirect incorporation of ^13^C, which are Nitrososphaeraceae, *Lysobacter* sp., and *Acidothermus* sp. (Figure 8). These ASVs are significantly different between H13 x L13 in the ANCOM analysis (Supplemental Figure S5), more abundant in H13 than L13, and show no change in the incubation with ^12^CH_4_ (H12 x L12), in addition to having a W-value greater than 10. The relative abundance of Nitrososphaeraceae ASVs is 5-fold higher in the ^13^C enriched region, with lower enrichment of *Lysobacter sp*. And *Acidothermus sp*. (Figure 8).

**Figure 8.**
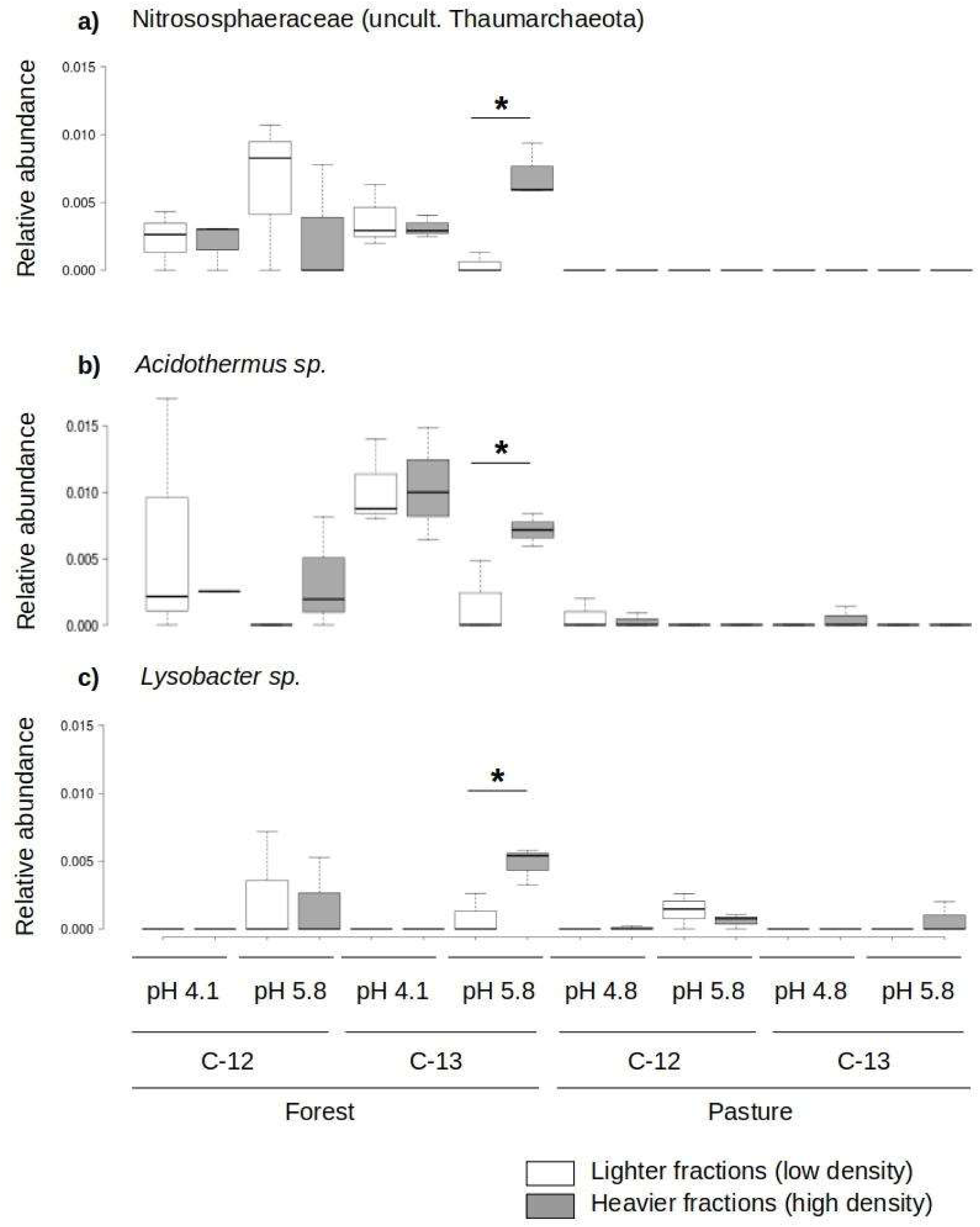
Relative abundance of ASVs from (a) Nitrososphaeraceae, (b) *Acidothermus* sp., and (c) *Lysobacter* sp. between heavy fractions (H) from the ^12^CH_4_ and ^13^CH_4_ incubations. *Significant differences (ANCOM).

## Discussion

The oxidation of methane in the soil can occur by high-affinity methanotrophs which operate at low methane concentrations, or by low-affinity methanotrophs adapted to high methane concentrations (Bender & Conrad, 1992). Since the rates of methane uptake in the soil can change depending on the concentration of methane to which soil is exposed, the availability of 10,000 ppm of methane applied in our experimental setup can be selected for a low-affinity response. In temperate forest soils, high-affinity methane consumption occurs up to 500 ppm, and low affinity can occur up to 30,000 ppm of methane in the atmosphere (Reay & Nedwell, 2004). The incubation with 10,000 ppm of ^13^CH_4_ showed that methane uptake in pasture soils can start not only quicker, but it also oxidizes methane at a rate five times higher than what was observed in forest soil, an indicator that communities in the two land uses may have a different methane consumption activity and/or composition.

Liming soils from both land uses led to greater methane uptake, which indicates the presence of a latent, low affinity, methanotrophic community, that can be stimulated by liming. The finding that liming promotes methanotrophy under low-affinity conditions in pasture soils suggests that pasture soils might have been pre-adapted to high methane concentrations. The effect of soil liming on methane cycling is poorly understood (Zhang et al., 2022). Liming tends to increase the oxidation of methane in temperate soils (Kunhikrishnan et al., 2016), but only a few studies are available on tropical soils, and results in the literature indicate no impact (Lammel et al., 2018; Mosier et al., 1998) or a reduction of methane consumption (Fonseca de Souza et al., 2022). Reports from temperate soils from both forest or agricultural origin describe either a decrease in methane consumption following liming (Borken et al., 2000; Butterbach-Bahl & Papen, 2002), or an increase (Barton et al., 2013; García-Marco et al., 2016; Hütsch et al., 1994).

Liming pasture soils leads to an increase in the methane absolute uptake 5-10 times higher than that of acidic forest soils. In temperate forest soils, the optimum of methane uptake under 100,000 ppm is observed at pH values close to 7.0 (Bender & Conrad, 1995), up to 3 times higher than under natural pH. These results lead us to expect that the tropical forest soils could have a similar response after liming, which was not observed. Using isotopic analysis of ^13^CO_2_, we estimated the ^13^C-incorporation into biomass. We found that up to 80% of ^13^C from the applied methane source (^13^CH_4_) was incorporated into the microbial biomass of pH-corrected pasture soils compared to an average of 50% incorporation in acidic pasture soils. During the same time interval acidic forest soils showed less than 5% incorporation, and liming increased biomass incorporation of ^13^C to only 10% in forest soil. Such results demonstrate the importance of soil pH as a modulator of methanotrophic activity in tropical soils, which had been acknowledged already for temperate forest soils (Amaral et al., 1998). In temperate forest soils, optimum methane uptake at atmospheric methane concentrations was found in the natural pH range, decreasing both with an increase or a decrease in soil pH (Amaral et al., 1998) and suggesting a soil microbial community well adapted to the natural pH value.

The abundance of active methanotrophs increased only in pasture soils, with and without liming. There was no detectable labeling of DNA belonging to methanotrophs in forest soils, which was expected given the low incorporation of ^13^C into the biomass in these soils and the absence of an increased H/L ratio of the *pmo*A gene. However, this evaluation of absolute abundance considered only those methanotrophs holding a *pmo*A gene, while other microorganisms capable of using atmospheric CH_4_ that do not have this gene, such as ammonia oxidizers, would not be detected (Bédard & Knowles, 1989). The reason for the non-enrichment of methanotrophs in forest soils may be due to the short incubation time in this experiment (Caro et al. 2022). In addition, due to soil homogenization by aggregate disruption and sieving, even formerly anoxic zones inside large soil aggregates are now exposed to atmospheric oxygen. This new availability of carbon stimulates the aerobic heterotrophic soil community, which can also affect methanotrophs. Using soils from the same region, without homogenization, Kroeger et al. (2021) carried out incubations with atmospheres at 30,000 ppm of ^13^CH_4_ over seven months, resulting in an enrichment of methanotrophs in forest soils. However, investigating forest soils from the western Amazon, the enrichment for isolation with minimal mineral medium under an atmosphere of 100,000 ppm ^13^CH_4_, Tessaro (2013) observed total uptake of methane after 60 hours of incubation, suggesting that different soils might result in very distinct methane uptake rates that are related to soil community composition and edaphic factors.

Members of two taxa of bacteria were identified as active in limed pasture soils. The first group consisted of two distinct ASVs and belonged to the family Beijerinckiaceae, a taxonomic level that may indicate the involvement of more than one genus or species. Both ASVs have a close phylogenetic relationship with methanotrophs known in this family and were detected in non-limed pasture soils and forest soils after liming, but in both instances without significant enrichment of heavy ^13^C. The second group consists of the genus *Methylocaldum* sp. which composes type I methanotrophs, has at least four isolated species, three from aquatic environments/sediments and one from agricultural soil, and is commonly detected in soils (Knief, 2015). Beijerinckiaceae are type II methanotrophs, with some exceptions in the genus *Beijerinckia* sp. This family includes a moderately acidophilic genus common in forest soils that clusters with ASV sequences found in our studies (Knief, 2015). Members of *Methylocella* sp., a genus within the Beijerinckiaceae, were detected in acidic pasture soils in the same region by Kroeger et al. (2021), but without observation of *Methylocaldum* sp. Tessaro (2013) detected *Methylocaldum sp*. in forest and pasture samples in the Western Amazon, but not Beijerinckiaceae. *Methylocaldum sp*. is a known member of soil microbial communities and a low-affinity methanotrophic genus, which exists in low abundance in acidic pasture soils and whose activity is increased after liming and exposure to high concentrations of methane. Our results are in accordance with what is known about type I methanotrophs showing low affinity, while type II methanotrophs are usually characterized by high affinity (below 40 ppm of methane) (Szafranek-Nakonieczna et al., 2019).

We could identify a significant increase in the relative abundance of three taxa within the dense ultracentrifugation fractions of the samples incubated with ^13^CH_4_. In contrast, the relative abundance of these three taxa did not change in the control incubation with ^12^CH_4_, which indicates active incorporation of ^13^C into their DNA in the experiment. The active groups were identified as Nitrososphaeraceae, *Lysobacter* sp., and *Acidothermus* sp. The first group is a family of archaea in the phylum *Crenarchaeota* and the class *Nitrososphaeria* (formerly *Thaumarchaeota*), associated with ammonia oxidation (Pester et al., 2011). The enzyme ammonia monooxygenase is capable of converting methane to methanol (Bédard & Knowles, 1989), and this may be the reason for its enrichment in the ^13^CH_4_ incubation. However, even if they are capable of converting methane to methanol, there is no record of the ability to use methanol by these microorganisms, or the presence of genes for methanol-dehydrogenase in available genomes. A syntrophic relationship is possible between Nitrososphaeraceae and *Lysobacter sp*., as available genomes of *Lysobacter sp*. indicate genes for methanol-dehydrogenase, and for the entire pathway to generate formate (Kruistum et al. 2018). Formate is an important regulator of cellular metabolism and is based on the synthesis of purines and other complex molecules (Brosnan & Brosnan, 2020). If secreted, formate could complex with cellulose (Fujimoto et al., 1986), and the resulting cellulose-formate would be a more easily absorbable molecule to be incorporated by *Acidothermus* sp., a taxon known for cellulolytic capacity (Mohagheghi et al., 1986).

## Conclusion

The liming of pasture soils in the Amazon region can significantly stimulate the uptake of methane under high methane concentrations within less than 30 days. However, conditions that create methane concentrations as high as 10,000 ppm in pasture soils are not well understood. In forest soils, acidity correction by liming has only a small stimulating effect on methane consumption over the same time interval, which indicates a microbial community adapted to low concentrations of methane. Methanotrophs from both metabolic groups of methane affinity were identified as active in limed pasture soils. The presence of active members of methanotroph type I, such as *Methylocaldum* sp., and type II, such as Beijerinckaceae indicated that both types of methanotrophs responded positively to liming in pasture soils and under conditions of low affinity. This work enables new questions about the potential of liming pastures in the Amazon region to mitigate soil methane emissions. Further field research is needed to decipher the dynamics of methanogenesis in these soils.

## Credit authorship contribution statement

**Leandro F. de Souza:** Conceptualization, Methodology, Formal analysis, Investigation, Data Curation, Project Administration, Writing - original draft, Writing - review & editing. **Fernanda M. Nakamura:** Methodology, Writing - review & editing. **Marie Kroeger:** Investigation, Writing - review & editing. **Dasiel Obregon:** Formal Analysis, Writing - review & editing. **Moacir T. Moraes:** Methodology, Writing - review & editing. **Mariana G. Vicente:** Methodology, Writing - review & editing. **Marcelo Z. Moreira:** Investigation, Methodology, Writing - review & editing. **Vivian H. Pellizari:** Investigation, Methodology. **Siu M. Tsai:** Conceptualization, Methodology, Resources, Funding acquisition, Writing - review & editing, Supervision. **Klaus Nüsslein:** Conceptualization, Methodology, Resources, Funding acquisition, Writing - review & editing, Supervision.

## Acknowledgments

The authors thank Mr. Bernardo for logistical support and permission to work on his land, and to the Large-Scale Biosphere-Atmosphere Program (LBA) coordinated by the National Institute for Amazon Research (INPA) for data access, logistical support, and infrastructure during fieldwork. Special thanks go to Prof. Plínio B. de Camargo and Wagner Piccinini for their invaluable assistance during field activities and to Prof. Luiz Domeignoz-Horta for his insightful suggestions, and critical reading of the draft.

## Supplementary Material

## Supplementary Figures and Methods

### Preliminary incubation (pilot experiment) under different concentrations of ^12^CH_4_

The efficiency of methane uptake in Amazon Forest and pasture soils under different atmospheric concentrations of this gas is an open question. To comprehend how methane uptake dynamics change in these soils, we set up a preliminary experiment exposing the soils in microcosms to different concentrations of methane in the head space. Microcosms were established with soils from the 2017 expedition, with 5 mm sieved soils from pooled sampling points, being assembled in gas tight flasks (volume 1.65 L) containing 400 g of soil (dry weight) forming a 10 cm layer. Microcosms with either Primary Forest or Pasture soil samples were incubated at 25°C, with five technical replicates each. The soil water content was kept constant at 70-80% of the water holding capacity. Artificial atmospheres were established by adding high purity ^12^CH_4_ gas (99.5% - White Martins, Piracicaba, SP, Brazil) with the aid of polypropylene syringes into the septum-sealed flasks, generating final methane concentrations of approximately 2 (negative controls, atmospheric rate), 200, 2,000 and 20,000 ppm of ^12^CH_4_, with concentration values verified by gas chromatography, as described below. Methane fluxes were assessed at weekly intervals, followed by replacement of the ^12^CH_4_ consumed.

We observed that 20% of the added methane was consumed over 7 days (Supplemental Figure S1). The ability to incorporate methane at concentrations of 200 ppm and 2,000 ppm was similar in both soils. In pasture soil, the average consumption rate of added methane under 200 or 2,000 ppm was the same, between 35% and 45%, doubling between days 17 and 24 (Supplemental Figure S1). In forest soils, methane consumption showed a tendency to increase by 100% to 200% after 7 days. The average methane consumption rate in the two soils was similar, and changed according to the initial concentrations except for the concentration of 20,000 ppm in forest soil, which remained low for 24 days.

**Supplemental Figure S1.**
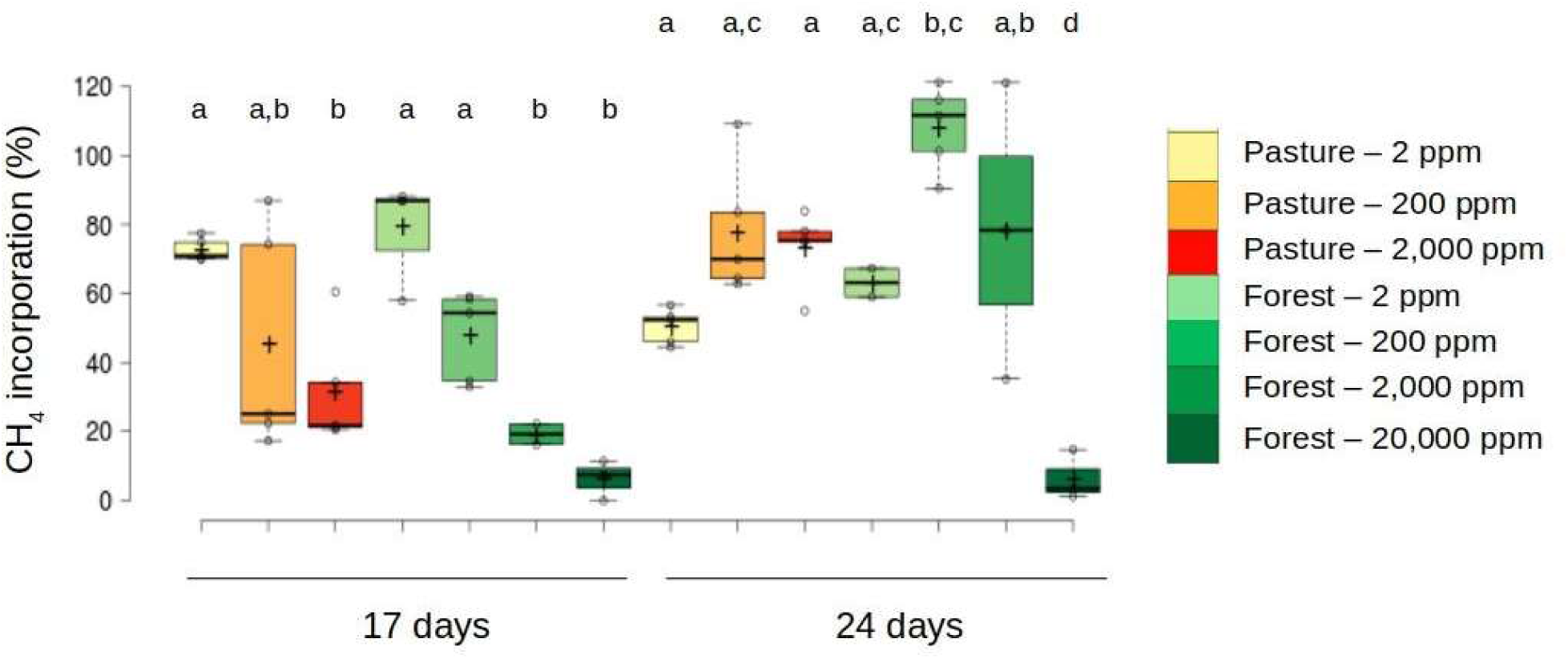
Percentage of ^12^CH_4_ oxidized in the microcosm under different methane concentrations in Pasture and Forest soils from Santarem, PA at four-day intervals. The mean values in the boxplots are indicated with a "+", and significant differences between groups are shown with letters above each boxplot for 17 and 24 days, respectively (Tukey HSD; p <0.05). P-2 ppm, P-2,000 ppm, FP-2 ppm and FP-200 ppm changed significantly between 17 and 24 days (Tukey HSD; p <0.05).

In this study, acidic rainforest soils were able to oxidize 80% of the applied methane under 2,000 ppm, in a four-day interval and after acclimatization for 24 days. Pasture soils performed similarly to forest soils under the same methane concentration. This indicates that the high affinity of tropical forest soils, already observed in Amazonian dark earth (Lima et al., 2014), can saturate at values greater than the 500 ppm of methane observed for temperate forest soils (Reay & Nedwell, 2004).

In the interval between 17 and 24 days of incubation, there was a reduction in the uptake capacity of the two land uses under 2 ppm. This can be due to starvation of methanotrophs, as observed in a study of temperate forest soil subjected to concentrations as low as 0.03 ppm of methane (Schnell & King, 1995). In our experiment, methane was replaced after four-day cycles, and was likely consumed prior to replenishment which exposed the community for some period of this four-day interval to concentrations lower than the atmospheric. Under 2,000 ppm incubation after 24 days, the uptake rate was similar to that observed after 17 days under 2 ppm in soils from both land uses. This differs from uptake rates observed in temperate forest soils incubated at 1,000 ppm, where the rate can be 100 times higher than under atmospheric concentrations (Schnell & King, 1995). When incubated under 20,000 ppm there is a significant decrease in the methane uptake in forest soil, which indicates that the microbial community responsible for the low affinity in the soil has little presence or activity in these conditions.

Ultracentrifugation followed by fractionation of the DNA extracted from these soils separated DNA from both incubations (^12^CH_4_ and ^13^CH_4_) into a low-density region, and a high-density and a very-high-density region (Supplemental Figure S2). Almost all DNA was found in the low-density region, and much less in the high-density regions. In incubations with high incorporation of ^13^C into the DNA of the community, a peak of DNA would be expected in the higher density regions of the samples incubated with ^13^C substrate, and no peak in samples incubated with ^12^C substrate. For example, a peak in the higher density fractions could be observed in incubations with substrates used by a large portion of the microbial community, such as ^13^C-glucose. But for substrates used by specialized guilds such as ^13^C-cellulose, the incorporation of ^13^C may not have labeled enough DNA for detection outside the fluorimetric error range. Results that illustrate this effect can be seen in López-Mondéjar et al. (2018).

Another indicator of fractionation quality is the distribution of DNA by GC composition of genomes between high- and low-density regions. The separation of environmental DNA by density gradient generates a gradient of GC/AT ratios and there is a tendency for genomes with higher GC content to be found in the dense region regardless of the incorporation of ^13^C isotopes (Dumont & Murrell, 2005). If the fractionation happened as expected, the GC ratio in the DNA sequences in the denser fractions is expected to be higher than in the light fractions. However, this effect should happen in a similar way in the ^12^C control, as the GC content in both situations would be equivalent. Thus, by comparative analysis of both incubations it is possible to exclude groups enriched by high GC content or other factors and not by incorporation of ^13^C. The results of this work showed a significant enrichment of microorganisms with high GC %mol content in the denser fractions.

**Supplemental Figure S2.**
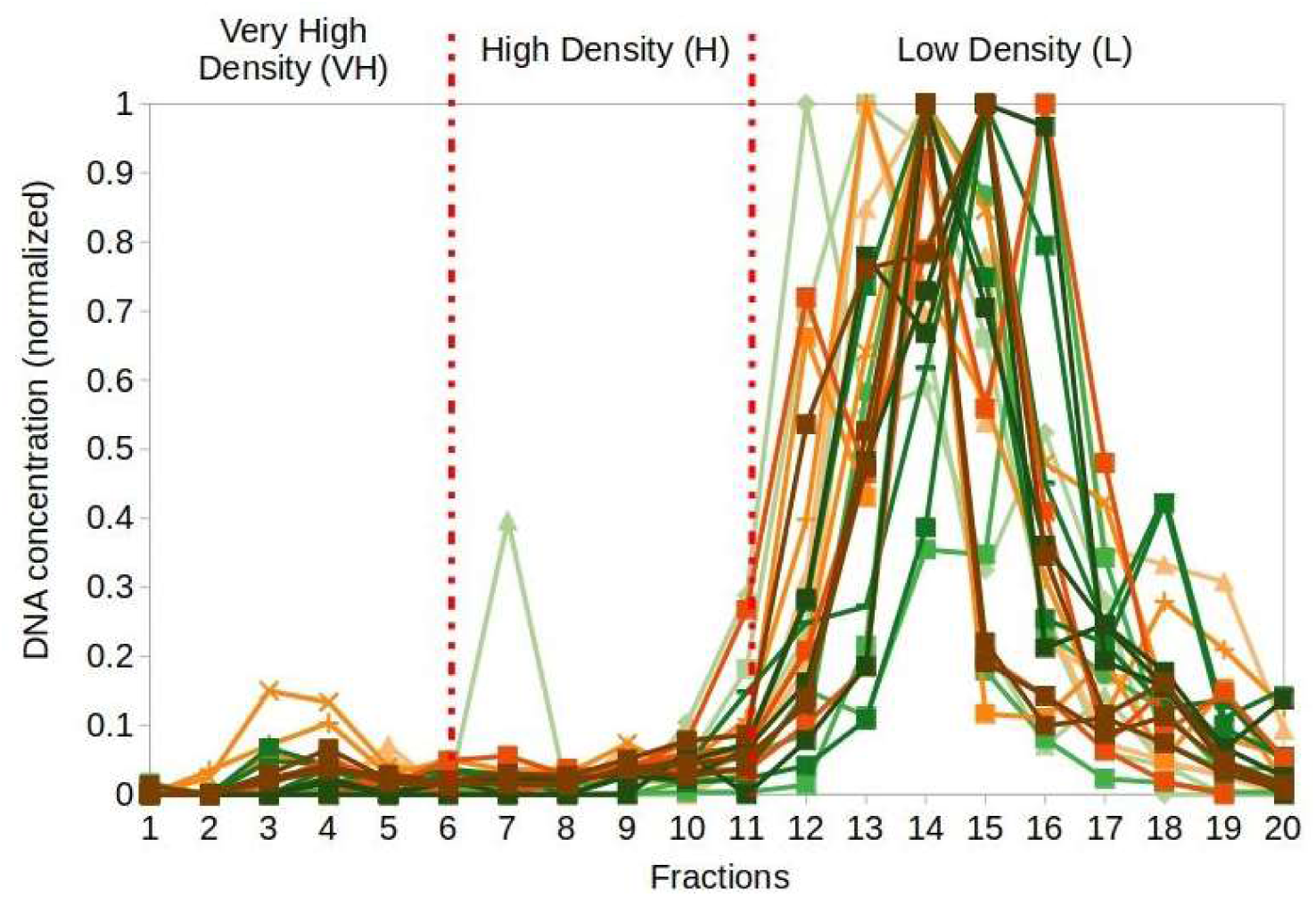
Normalized DNA concentration (ratio) in relation to the highest concentration observed in each sample after fractionation. Regions grouped by density are defined as: VH (fractions 1-6), H (7-11), and L (12-20). Orange hues represent pasture samples and forest samples are identified by green hues.

**Supplemental Figure S3:**
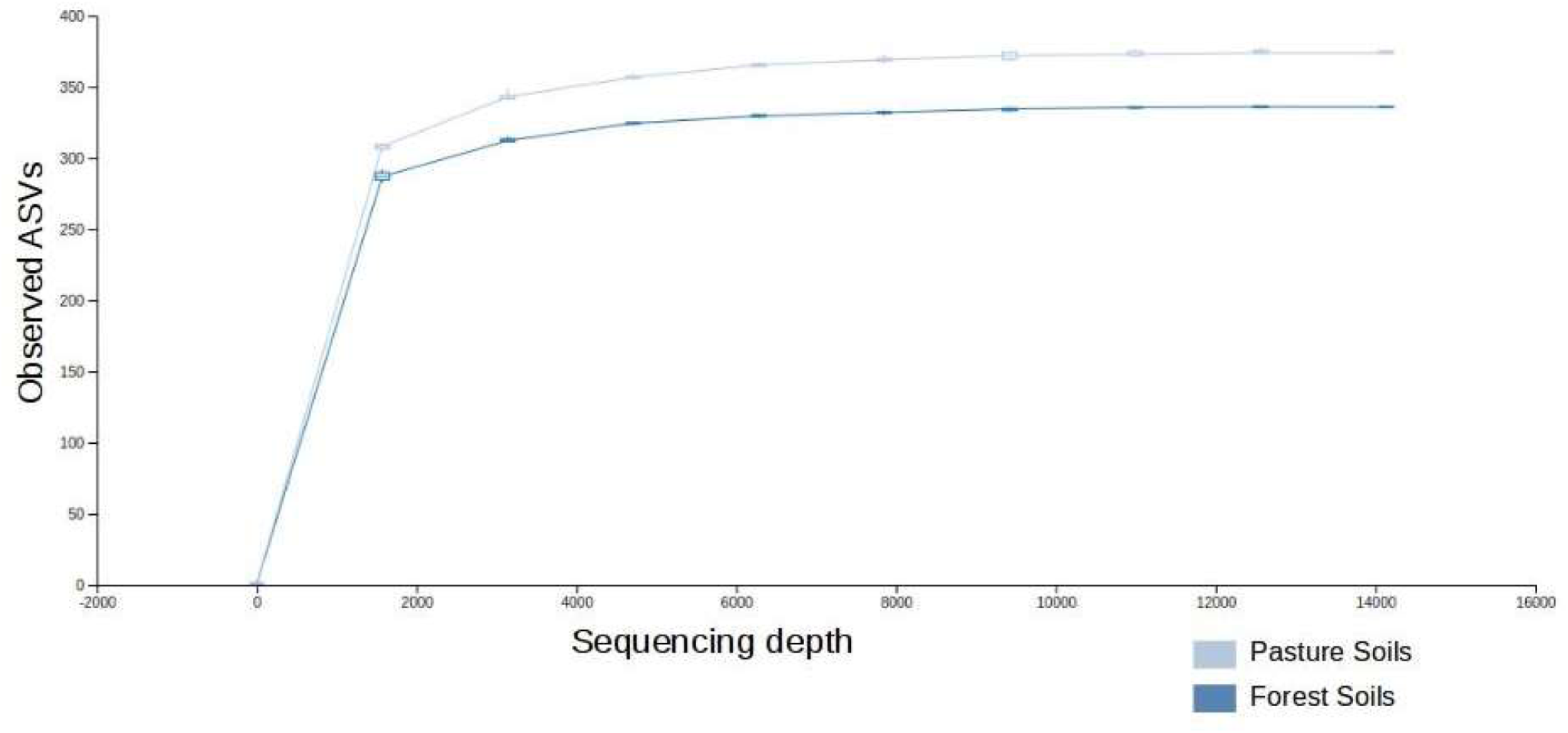
Rarefaction plots of ASVs identified in DNA from both soil treatments after SIP analysis.

**Supplemental Figure S4.**
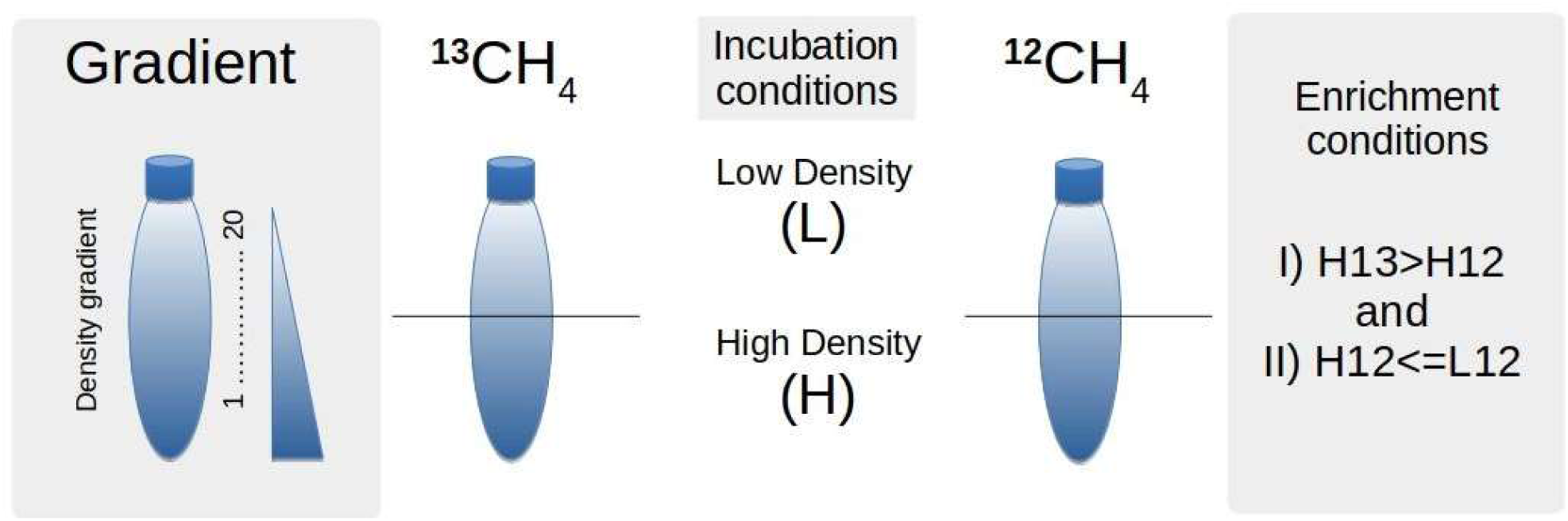
Schematic representation of the conditions established *a priori* for the fractionation using DNA-SIP. The relative abundance of ASVs from different gradient points (low or high density) was assessed according to these criteria, which allowed us to qualify if DNA sequences were enriched in ^13^C and, therefore, indicative of ^13^C assimilation activity. The conditions were I) ASVs should be enriched in the high-density region of the ^13^CH^4^ incubation rather than in the high-density region of the ^12^CH_4_ incubation. This allows us to detect microorganisms that incorporated ^13^C, despite their buoyant density related to their DNA %GC content; II) ASVs should not be enriched within the high-density region of the ^12^CH_4_ incubation compared to the low-density region.

**Supplemental Figure S5:**
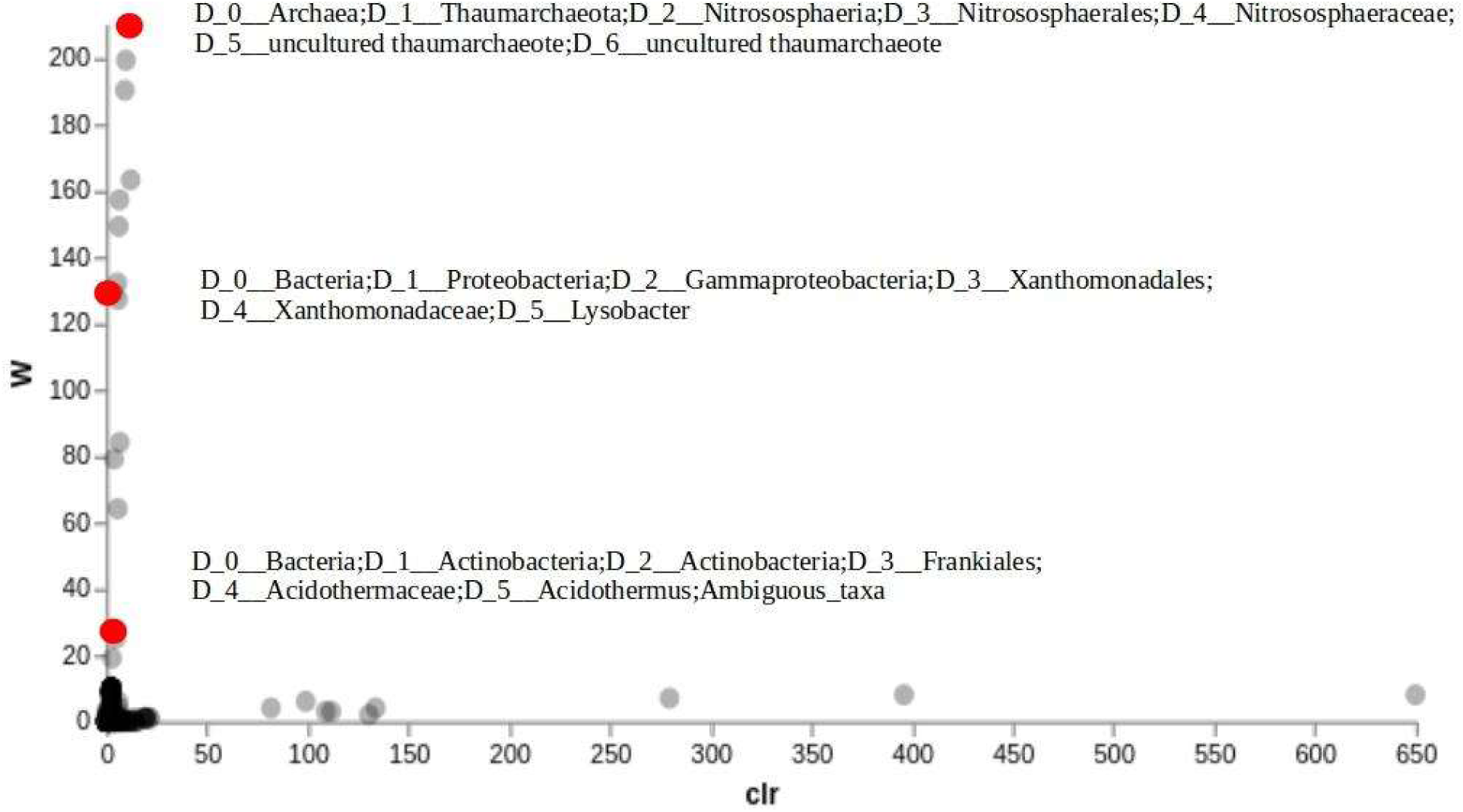
Consistent enrichment of certain ASVs after incubation of limed forest soil with ^13^CH_4_. Results of an ANCOM analysis, comparing high-density DNA (H13) with low density DNA (L13). Centered Log Ratio (CLR) indicates how much a group is enriched in relation to the average relative abundance of a respective ASV, and W indicates the consistency of this observation in relation to the other ASVs. The higher the values of W and CLR are, the greater and more consistent is the difference in relative abundance. Red circles highlight significantly different ASVs, limited by the following characteristics: W> 10, no differences between L12xH12 and higher relative abundance in H13 than in L13.

**Table S1:** Physicochemical characteristics of sampled forest and pasture soils.

| Property | Unit | F-natural pH | F-limed | P-natural pH | P-limed |
| --- | --- | --- | --- | --- | --- |
| pH | CaCl<sub>2</sub> | 3.7 ± 0.1 | 5.3 ± 0.2 | 4.1 ± 0.1 | 5.0 ± 0.2 |
| pH | H<sub>2</sub>O | 4.1 ± 0.2 | 5.8 ± 0.2 | 4.8 ± 0.1 | 5.6 ± 0.1 |
| O.M. | g.Kg<sup>-1</sup> | 40.8 ± 7.2 | 34.8 ± 3.1 | 28.4 ± 5.9 | 27.0 ± 4.6 |
| N | mg.Kg<sup>-1</sup> | 2156 ± 656 | 1960 ± 531 | 1260 ± 131 | 1330 ± 221 |
| K |  | 1.0 ± 0.1 | 1.0 ± 0.1 | 1.0 ± 0.1 | 1.0 ± 0.1 |
| Ca |  | 3.4 ± 0.5 | 53.2 ± 3.6 | 7.2 ± 0.8 | 30.8 ± 1.5 |
| Mg |  | 2.0 ± 0 | 3.0 ± 0 | 4.6 ± 0.9 | 4.0 ± 0 |
| H+Al | --cmolc.dm<sup>-3</sup>-- | 127.6 ± 18 | 33.0 ± 3.7 | 64.4 ± 5.0 | 34.2 ± 2.5 |
| Al |  | 18.6 ± 1.1 | 0.8 ± 0.8 | 7.4 ± 1.1 | 1.0 ± 0.7 |
| BS |  | 6.4 ± 0.5 | 57.2 ± 3.6 | 12.8 ± 1.5 | 35.8 ± 1.5 |
| CEC |  | 134 ± 18.3 | 90.2 ± 1.6 | 77.2 ± 4.8 | 70.0 ± 2.9 |
| Cu |  | 0.4 ± 0 | 0.3 ± 0.1 | 0.3 ± 0.1 | 0.3 ± 0.2 |
| Fe |  | 146 ± 41.3 | 81.8 ± 19.6 | 138.8 ± 7.0 | 80.4 ± 15.2 |
| Zn |  | 36.4 ± 27.1 | 38.4 ± 18.3 | 30.8 ± 11.0 | 39.8 ± 17.7 |
| Mn | --mg.dm<sup>-3</sup>-- | 10.4 ± 1.1 | 6.5 ± 2.0 | 6.3 ± 1.0 | 4.9 ± 1.3 |
| B |  | 0.8 ± 0.2 | 0.8 ± 0.2 | 0.4 ± 0.1 | 0.5 ± 0.2 |
| P |  | 5.0 ± 0.7 | 4.2 ± 0.4 | 2.6 ± 0.5 | 3.0 ± 0.7 |
| S |  | 18.6 ± 7.3 | 16.2 ± 2.7 | 9.6 ± 1.3 | 9.4 ± 0.5 |
| V | % | 4.8 ± 0.4 | 63.6 ± 4.2 | 16.8 ± 2.3 | 51.4 ± 2.1 |
| m | % | 74.4 ± 2.6 | 1.4 ± 1.3 | 36.8 ± 6.0 | 2.8 ± 1.8 |
| Clay |  | 817.6 ± 54.6 | 801.6 ± 10.5 | 489 ± 18.8 | 487.2 ± 30.9 |
| Silt |  | 122.4 ± 51.4 | 94.4 ± 36.8 | 47.0 ± 43.6 | 50.8 ± 14.5 |
| Total Sand | -- g.kg<sup>-1</sup> -- | 60 ± 18.7 | 104 ± 37.8 | 464 ± 21.7 | 462 ± 17.9 |
| Thick Sand |  | 40 ± 12.2 | 70 ± 29.2 | 318 ± 32.7 | 314 ± 28.8 |
| Thin Sand |  | 20 ± 7.1 | 34 ± 11.4 | 146 ± 11.4 | 148 ± 13.0 |

F = Forest; P = Pasture; O.M. = Organic Matter; BS = Base Summary; CEC = Cation Exchange Capacity; V= Base saturation; m = Aluminum saturation; H+ Al = Potential acidity.

**Table S2:** Sequencing depth of 16S *rRNA* amplicons after filtering, denoising, merging forward and reverse sequences, and removal of chimeras by the DADA2 software package.

| Incubation | Treatment | Number of ASVs |
| --- | --- | --- |
| <sup>12</sup>CH<sub>4</sub> | F-natural pH-H | 32780 ± 13493 |
|  | F-natural pH-L | 40950 ± 8227 |
|  | P-natural pH-H | 37911 ± 17888 |
|  | P-natural pH-L | 29913 ± 5201 |
|  | P-natural pH-VH | 41184 ± 22041 |
|  | F-liming-H | 35068 ± 11969 |
|  | F-liming-L | 25364 ± 4256 |
|  | P-liming-H | 36848 ± 14201 |
|  | P-liming-L | 33737 ± 4788 |
|  | P-liming-VH | 38985 ± 9222 |
| <sup>13</sup>CH<sub>4</sub> | F-natural pH-H | 37573 ± 7974 |
|  | F-natural pH-L | 43027 ± 8995 |
|  | P-natural pH-H | 32371 ± 3593 |
|  | P-natural pH-L | 24589 ± 2449 |
|  | P-natural pH-VH | 33050 ± 7030 |
|  | F-liming-H | 44402 ± 12225 |
|  | F-liming-L | 34880 ± 28860 |
|  | P-liming-H | 31821 ± 12540 |
|  | P-liming-L | 24483 ± 17482 |
|  | P-liming-VH | 49749 ± 44474 |

F = Forest; P = Pasture. H = High Density; L = Low Density; VH = Very High Density.

